# *In Silico* Targeting of Calpain-5 in Autosomal Dominant Neovascular Inflammatory Vitreoretinopathy (ADNIV) with Peptides and Natural Products

**DOI:** 10.64898/2026.08.11.744189

**Authors:** Surya Charan Garimella, Yash Bhargava

## Abstract

Autosomal dominant neovascular inflammatory vitreoretinopathy (ADNIV) is a rare retinal disease caused by gain-of-function mutations in the non-classical calcium-activated cysteine protease calpain-5 (CAPN5). These mutations lower the calcium threshold for catalytic-triad alignment with downstream effects including excessive proteolysis and retinal degeneration, making CAPN5 a therapeutic target. Clinical studies showed that knockout of calpain-5 resulted in no negative side effects, supporting therapy through inhibition. We mapped the druggable pockets of CAPN5 with a 500 ns phenol cosolvent molecular dynamics (MD) simulation. Occupancy analysis resolved five pockets, against which 448,314 COCONUT natural products were screened with Uni-Dock (2,241,570 docked combinations). In parallel, BoltzGen was used to design peptide binders against multiple candidate regions, from which three were selected: the PC1–PC2 subdomain interface, the PC2 regulatory loop (PC2L1) and the catalytic region. The top three designs were co-folded with Boltz-2 at high interface confidence (ipTM 0.91–0.95). The top three peptides and four small molecules were then simulated against wild-type CAPN5 and the four canonical ADNIV variants R243L, L244P, K250N and R289W, each condition in independent triplicate, giving 105 production simulations of 100 ns. Scoring by MM-PBSA revealed favorable peptide interface energies, the most favorable being the largest of the three designs (ΔTOTAL −58.6 ± 5.6 kcal/mol for a 23-residue peptide against wild type), while the small-molecule panel returned −8.7 to −23.8 kcal/mol. A total of 15.8 *µ*s of cosolvent, filtering, and production MD prioritizes the catalytic cleft and an adjacent groove for experimental testing and provides candidate peptide and small-molecule binders for evaluating CAPN5 inhibition in ADNIV.

## 1 Introduction

Autosomal dominant neovascular inflammatory vitreoretinopathy (ADNIV) is a rare inherited retinal disease in which intraocular inflammation, retinal neovascularization, and progressive photoreceptor degeneration accumulate over decades [1, 2, 3]. The disease passes through five progressive phases beginning in adolescence (early inflammation, retinal degeneration, neovascularization, fibrosis and end-stage disease), with severe visual loss typically occurring between the fourth and fifth decades. Late-stage complications include tractional retinal detachment, neovascular glaucoma, and phthisis bulbi.

Current therapeutic efforts target the downstream inflammatory, neovascular, and fibrotic consequences of the disease. Corticosteroids, including sustained-release fluocinolone acetonide implants, anti-VEGF (anti-vascular endothelial growth factor) injection, immunosuppression, and laser photocoagulation provide temporary control of symptoms, but none actually halts progression and several introduce iatrogenic burdens of their own [4, 5, 6, 7]. These side effects include steroid-induced ocular hypertension, glaucoma, cataract, and fibrotic encapsulation of the implant, all of which conflict with the goals of therapy.

The *CAPN5* gene encodes calpain-5, a non-classical member of the calpain family of calcium-activated cysteine proteases. It includes an N-terminal anchor region, a protease core with the Cys–His–Asn catalytic triad, and regulatory domain III and C2 regions. Wild-type calpain-5 performs limited proteolysis, stably cleaving products with neo-N-termini [8]. The proper Ca^2+^ threshold activation is required to follow through with these functions [9].

The ADNIV-associated substitutions R243L, L244P, K250N and R289W are the best-characterized pathogenic variants [1, 9] and cluster on flexible loop structures that restrain access to the catalytic site. Mutations lower the calcium threshold for activation, resulting in ADNIV disease progression [10]. Rarer variants such as G267S have been reported in only one or two patients and remain of uncertain pathogenicity, so were excluded from this study. All candidate inhibitors were evaluated against each canonical disease-associated mutant. Inhibition has been supported by a variety of findings to be the appropriate therapeutic logic for this target. Photoreceptor-specific *Capn5* knockout in mice preserves normal retinal structure and function, indicating that loss of calpain-5 activity is well tolerated. That study also identified 22 human *CAPN5* loss-of-function variants across 60,706 unrelated subjects without severe disease, and concluded that localized CAPN5 inhibition is a viable strategy against hyperactivating alleles [11]. Complementary work showed that lentiviral delivery of a human gain-of-function *CAPN5* variant into mouse retina is sufficient to reproduce ADNIV features [12], and that mutant *CAPN5* expression can be suppressed without cytotoxicity by RNA interference [13]. These studies support the inhibition of mutant protease activity as a therapeutic strategy. Selectivity is the primary challenge in targeting CAPN5. The classical calpains CAPN1 and CAPN2 are essential [14, 15, 16, 17, 18], so a pan-calpain inhibitor would be unusable as a chronic intravitreal therapy. Thus, distinct areas of the CAPN5 protease, such as the PC1–PC2 interface and regulatory loops, are better targets than the catalytic triad common to many proteases.

The surface of CAPN5 was probed using cosolvent MD with phenol-containing solvent to identify possible drug-binding pockets [19, 20]. These sites were used for both high-throughput natural-product screening and diffusion-based generative peptide design. Prioritized binders of both modalities were evaluated by explicit-solvent MD and end-point binding free-energy calculation against wild-type CAPN5 and each of the four canonical ADNIV variants.

## 2 Materials and Methods

### 2.1 Assembly of a Full-Length CAPN5 Structural Model

The experimental structure of the CAPN5 protease core was retrieved from the Protein Data Bank (PDB 6P3Q; 2.8 Å) [9, 21], which contains two identical chains spanning residues 4–349 and resolves the protease core (PC) region. An AlphaFold model of full-length human CAPN5 was retrieved from the AlphaFold Protein Structure Database [22]. The two models were superposed over their common ordered region (mean absolute error *<* 0.4 Å) and grafted to produce the final construct. The assembled model was energy-minimized by steepest descent to a maximum force of 500 kJ mol^−1^ nm^−1^ with particle-mesh Ewald electrostatics and a Verlet cutoff scheme. The resulting structure was used for all production MD and was used as the docking receptor after protonation at pH 7.4.

The catalytic triad is Cys81, His252 and Asn284, with Cys81 in protease core subdomain PC1 and His252 and Asn284 in PC2, so that a site spanning the two subdomains must contact residues on both sides of that division.

### 2.2 Modeling of ADNIV Variants

Four clinically reported ADNIV substitutions (R243L, L244P, K250N and R289W) were modeled onto the full-length construct. The variants are predicted to weaken the gating loop through distinct mechanisms including loss of electrostatic interactions (R243L), altered secondary-structure constraints (L244P), disruption of local hydrogen bonding (K250N), and steric effects introduced by R289W.

### 2.3 Phenol Cosolvent MD

To identify candidate druggable hotspots and cryptic pockets, a phenol cosolvent simulation was performed in OpenMM [23] following the methodology outlined by MixMD-style cosolvent simulations [19, 20]. The protein was parameterized with AMBER14/ff14SB [24], water with TIP3P-FB [25], and phenol probes with the Open Force Field 2.2.1 (Sage) small-molecule force field [26]. The system was solvated in a dodecahedral box with 1.2 nm padding and 0.15 M NaCl, and phenol molecules were added to a final concentration of 0.50 M, matching the probe concentration used for cryptic-pocket cosolvent simulations in CosolvKit [27]. Phenol was selected due to its aromatic hydrophobic face and hydrogen-bond donor/acceptor hydroxyl that report on both apolar and polar ligand-binding propensity. Hydrogen mass repartitioning to 3.024 amu was applied to allow a 4 fs integration timestep. Following energy minimization, the system was equilibrated through sequential NVT and NPT stages at 300 K and 1 bar. The 500 ns production simulation used a Langevin middle integrator for temperature control at 300 K, particle-mesh Ewald long-range electrostatics and a 1.0 nm nonbonded cutoff, with coordinates saved every 100 ps.

### 2.4 Hotspot Identification and Pocket Definition

Production trajectories were aligned to the initial frame on protein backbone atoms, and phenol positions were extracted. Occupancy was accumulated on a 0.5 Å grid, and cells with *>*10% of the maximum observed density within 7.0 Å of protein heavy atoms were clustered with DBSCAN (*ε* = 3.0 Å, minimum samples = 10) [28]. Hotspots were then ranked by total phenol observations and residue contacts were assigned when phenol heavy atoms fell within 4.5 Å of protein heavy atoms. These candidate pockets were then used for ligand docking and simulations. Occupancy scores were written to the B-factor column of a PDB file and rendered on the molecular surface in PyMOL [29].

### 2.5 High-Throughput Molecular Docking

The COCONUT natural product library was used for its size and coverage of drug-like natural chemical space [30]. Records were converted from SDF to PDBQT with Meeko, and entries that failed preparation or violated the docking constraints were removed. Next, ligands were filtered to ≤ 48 rotatable torsions and maximum dimensions of 16.5 Å, giving a final library with 448,314 compounds.

The prepared library was docked into each of the five cosolvent-derived hotspots on the pH 7.4 receptor using Uni-Dock with the Vina scoring function [31], whose GPU implementation enables large-scale screening. Cubic boxes of 15 × 15 × 15 Å were centered on the five hotspot centers transformed into the receptor coordinate frame. Docking using Uni-Dock’s fast search mode retained a single best-scoring pose per ligand per site, trading exhaustiveness of the conformational search for the throughput needed at this library size. The complete docking consisted of 448,314 ligands × 5 sites = 2,241,570 docked combinations.

### 2.6 Physicochemical Triage of Prioritized Compounds

Physicochemical, drug-likeness, and predicted-ADMET filters were used to remove compounds likely to be toxic or incompatible. Predicted absorption, distribution, metabolism, excretion and toxicity properties were computed with ADMET-AI [32], which reports both absolute predictions and percentile rankings against an FDA drug reference set. Lipophilicity was constrained by a soft LogP window of 0.8–3.8 to exclude compounds likely to suffer permeability. Due to bioavailability concerns, compounds below the 25th percentile for aqueous solubility were removed, and predicted genotoxicity was filtered at AMES *>* 0.6.

### 2.7 Diffusion-Based Peptide Binder Design

BoltzGen, an all-atom generative model for *de novo* binder design, was used for peptide design through refining noisy coordinates into binder–target complexes using a diffusion process [33]. Multiple candidate regions were specified from the structural and cosolvent analysis, and three were carried forward: the PC1–PC2 subdomain interface, which carries the loops that control subdomain alignment; the PC2 regulatory loop (PC2L1); and the catalytic region. Designs against the remaining regions scored less favorably at co-folding and were not advanced. Target residues were selected manually with the top 20 binders saved per site. Designed peptides were co-folded with CAPN5 using Boltz-2 [34] with the 6P3Q protease-core template. Interface confidence was assessed from ipTM, pTM, complex pLDDT and interface predicted aligned error, and the buried interface area and interface residue contacts were measured directly from the co-folded models.

### 2.8 MD-Based Peptide Filtration

Designed peptides were filtered with 160 short MD simulations. Conformational stability assessment occurred through 30 ns simulations (2 fs timestep, no hydrogen mass repartitioning) and scored by per-residue root-mean-square fluctuation (RMSF). The three lowest RMSF candidates per site were advanced to 100 ns simulations, from which MM-PBSA binding free energies were computed over the final 20 ns; the single strongest binder per site was retained. The three retained designs were then simulated against wild-type CAPN5 and each ADNIV variant.

### 2.9 MD Simulation Protocol

Production simulations were performed with GROMACS 2025.3 [35] using the CHARMM36 force field (July 2022 revision) [36] with hydrogen mass repartitioning to 3.024 amu, enabling a 4 fs timestep [37]; runs rebuilt to test integrator sensitivity were performed at 2 fs without repartitioning. Systems were solvated in dodecahedral boxes of TIP3P water [38] with a minimum 1 nm solute boundary separation, and neutralized with Na^+^, Cl^−^ and Ca^2+^ at 0.15 M NaCl and 0.1 M CaCl_2_, chosen to match conditions used in prior CAPN5 conformational studies.

Energy minimization used steepest descent to 500 kJ mol^−1^ nm^−1^ with particle-mesh Ewald electrostatics [39], a 1.2 nm real-space cutoff and force-switched van der Waals interactions between 1.0 and 1.2 nm under a Verlet scheme with a 0.1 nm buffer. Hydrogen bond constraints were applied during minimization and all-bond constraints thereafter using LINCS [40]. NVT and NPT equilibration were each run for 100 ps at the production temperature with a 2 fs timestep, with velocities initialized from a Maxwell–Boltzmann distribution, pressure held at 1 bar with a C-rescale barostat under isotropic coupling and temperature maintained with a velocity-rescaling thermostat [41]. Production runs were carried out at 310 K for 100 ns, using LINCS, with pressure controlled by a Parrinello–Rahman barostat [42]. Every binder–background condition was run in independent triplicate rather than once. Seven prioritized binders across five backgrounds at three replicates gives 105 production simulations of 100 ns (Table S3 of the Supplementary Material).

Each variant was simulated apo and in complex with prioritized binders under this protocol, providing the control against which ligand efficacy was assessed.

### 2.10 Binding Free Energy and Per-Residue Decomposition

End-point binding free energies were computed with gmx MMPBSA 1.6.4 [43] using the single-trajectory protocol at 298.15 K, decomposing the binding free energy as

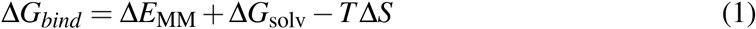

where Δ*E*_MM_ is the gas-phase molecular-mechanics interaction energy (electrostatic plus van der Waals), Δ*G*_solv_ the solvation free energy, and −*T* Δ*S* the entropic term. AMBER topologies were generated with ff99SB for protein [44] and GAFF for ligands [45]. The solvation free energy was split into polar and nonpolar components,

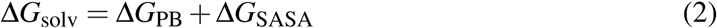

with the polar term Δ*G*_PB_ evaluated by the Poisson–Boltzmann model using solute and solvent dielectric constants of 1 and 80 respectively and the nonpolar term Δ*G*_SASA_ obtained from the solvent-accessible surface area; generalized-Born energies were computed in parallel for comparison. The entropic term was estimated by the interaction-entropy method over the final 25% of each trajectory,

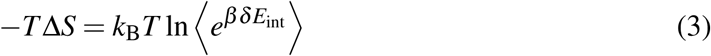

where *δ E*_int_ is the fluctuation of the gas-phase interaction energy about its mean and *β* = 1*/k*_B_*T*. Per-residue energy decomposition was performed to identify the residues contributing most to binding. Per-residue C*α* root-mean-square fluctuation, complex backbone root-mean-square deviation and radius of gyration were computed with the GROMACS analysis tools after least-squares fitting on the protein backbone.

Every energy reported below is the mean over the replicates of a condition, and every stated uncertainty is the standard deviation across those replicates rather than the inter-snapshot spread inside one trajectory.

## 3 Results and Discussion

### 3.1 A Full-Length CAPN5 Construct Was Used for Simulation

In the absence of an experimental structure of full-length CAPN5, a full-length model was assembled. Superposition of the AlphaFold model onto the 6P3Q protease-core crystal structure over their shared ordered region and using it as the scaffold while grafting the unresolved domain III, C2 and tail regions from the prediction produced a continuous structure for simulation (Fig. 1A). The complete pipeline built on this construct is summarized in Fig. 1C.

**Figure 1.**
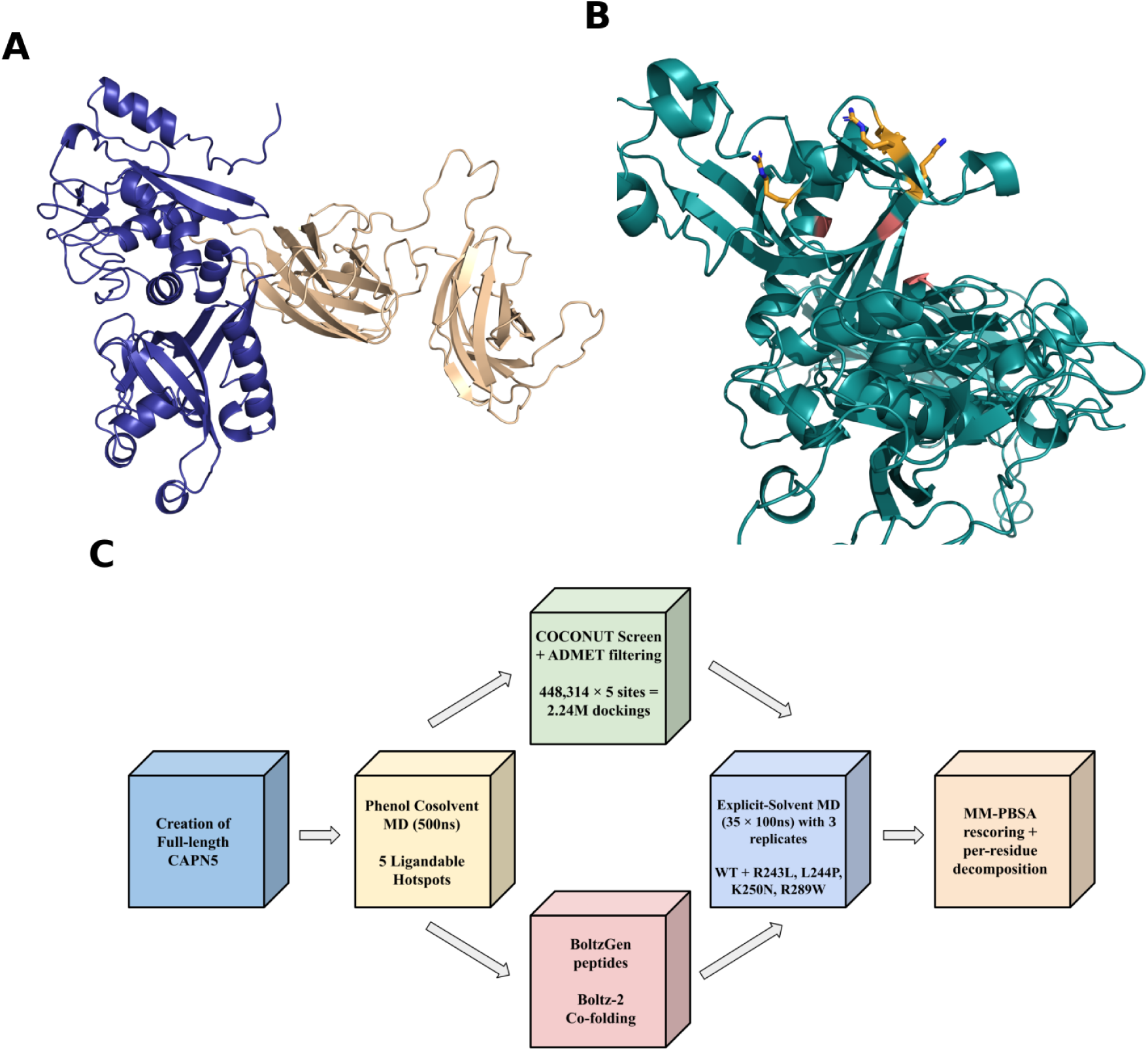
Full-length CAPN5 model and computational workflow. (A) The full-length CAPN5 construct used throughout the study, combining the experimental protease core (PDB 6P3Q, 2.8 Å ; blue) with AlphaFold-derived domain III, C2 and C-terminal tail regions grafted after superposition (wheat; MAE *<* 0.4 Å). (B) The four modeled ADNIV substitutions (R243L, L244P, K250N, R289W; sticks colored by atom, carbons orange) mapped onto the construct. (C) Overview of the workflow.

### 3.2 Phenol Cosolvent MD Resolves Five Ligandable Hotspots on CAPN5

The druggable pockets of CAPN5 were mapped using a 500 ns phenol cosolvent simulation at 0.50 M probe concentration (4 fs with hydrogen mass repartitioning). The occupancy clustering resolved five discrete hotspots which screening and design was then conducted against (Fig. 2; Table 1).

**Figure 2.**
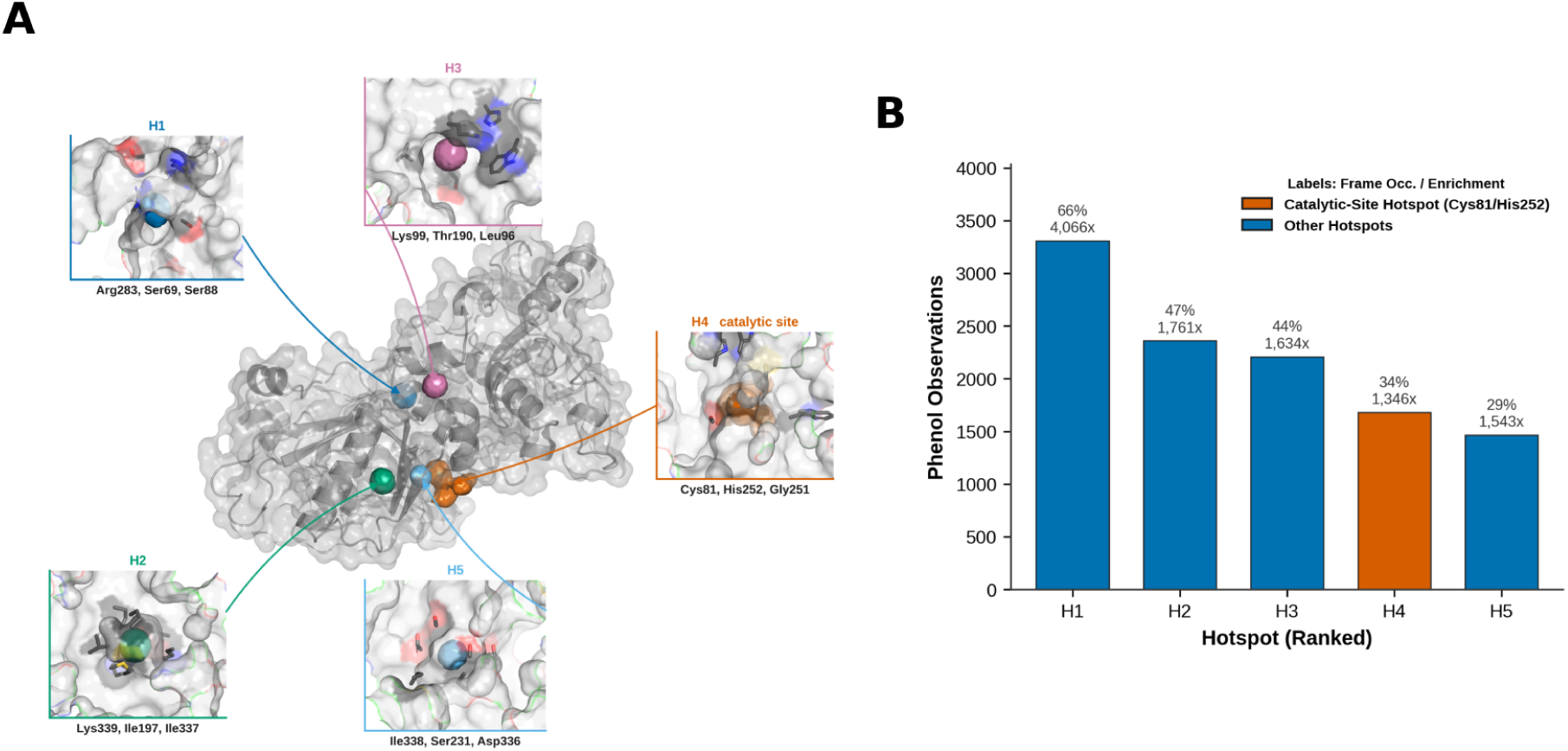
Phenol cosolvent molecular dynamics resolves five druggable hotspots. (A) The five phenol occupancy hotspots on the full-length CAPN5 surface, with a zoomed view of every pocket joined to its own density by a color-matched arrow and labeled with its top three contact residues (full contact sets in Table 1). (B) The five DBSCAN-resolved hotspot clusters ranked by phenol observations; hotspot 4 (catalytic) is highlighted.

**Table 1.** Cosolvent Hotspots on CAPN5. Phenol-enriched hotspots resolved by DBSCAN clustering of cosolvent occupancy of the 500 ns simulation, catalytic residues bolded.

| Rank | Phenol obs. | Frame occ. (%) | Max-cell enrich. (×) | Dist. to protein ($\text{\AA}$) | Principal contact residues |
| --- | --- | --- | --- | --- | --- |
| 1 | 3307 | 66.1 | 4066 | 1.9 | Arg283, Ser69, Ser88, His71, Pro285, Asp72, Leu73, Ser92, Ala255 |
| 2 | 2359 | 47.2 | 1761 | 0.8 | Lys339, Ile197, Ile337, Leu214, Met218, Arg217, Glu195, Ile338 |
| 3 | 2208 | 44.2 | 1634 | 1.8 | Lys99, Thr190, Leu96, Phe189, Trp131, Val100, Leu90, His124 |
| 4 | 1680 | 33.6 | 1346 | 1.7 | Gly251, Ala253, **Cys81**, **His252**, Ser231, Ala85, Trp82, Trp286, Thr182 |
| 5 | 1465 | 29.3 | 1543 | 2.0 | Ile338, Ser231, Asp336, Thr182, Ser229, Ile337, Ala230, Ala253, Pro196 |

The leading hotspot accumulated 3307 phenol observations, appeared in 66.1% of analyzed frames, and reached 4066-fold maximum-cell enrichment relative to bulk phenol concentration, contacting Arg283, Ser69, Ser88, His71, Pro285, Asp72, Leu73, Ser92, and Ala255. The remaining four accumulated 1465 to 2359 observations at frame occupancies between 29.3% and 47.2%, with enrichments of 1346- to 1761-fold (Fig. 2B).

The fourth-ranked hotspot contacts Cys81 and His252, two members of the catalytic triad, along with Gly251, Ala253, Ser231, Trp82 and Trp286, and therefore shows the catalytic site as a solvent-accessible, probe-enriched region. Hotspots 2 and 5 share Ile337 and Ile338 among their top-ranked contacts. Hotspot 5 overlaps more with the catalytic hotspot than with hotspot 2, sharing Ser231 and Ala253 among its top-ranked contacts and Thr182 and Ala85 below that cut. Hotspot 2 is a separate pocket lying entirely within residues 195–339, with no contact on the Cys81 side of the protease core, and so does not span the two subdomains. The hydrophobic probe therefore resolves the catalytic cleft and the adjacent groove of hotspot 5 as the two druggable surfaces the mapping supports, both of them on the catalytic face of the protease core rather than at a subdomain interface.

### 3.3 High-Throughput Docking of the COCONUT Library Ranks Natural Products Across the Five Hotspots

After filtering for ≤48 torsions and ≤16.5 Å maximum dimension, 448,314 prepared CO-CONUT compounds were docked against the five hotspots, yielding 2,143,693 Uni-Dock scores (Fig. 3A). Score distributions differed sharply between pockets (Fig. 3B,C; Table 2). Site 4 and site 5 were broadly accessible, returning 261,287 and 150,156 compounds at or below −6 kcal/mol respectively, with the best scores of −11.02 and −10.54 kcal/mol. Sites 2 and 3 were intermediate (7311 and 46,273 compounds below −6 kcal/mol; best scores −9.01 and −9.31 kcal/mol). Site 1 was unfavorable for docking, with only 32 of 425,475 compounds scoring ≤ −6 kcal/mol and a positive median score of +13.98 kcal/mol.

**Figure 3.**
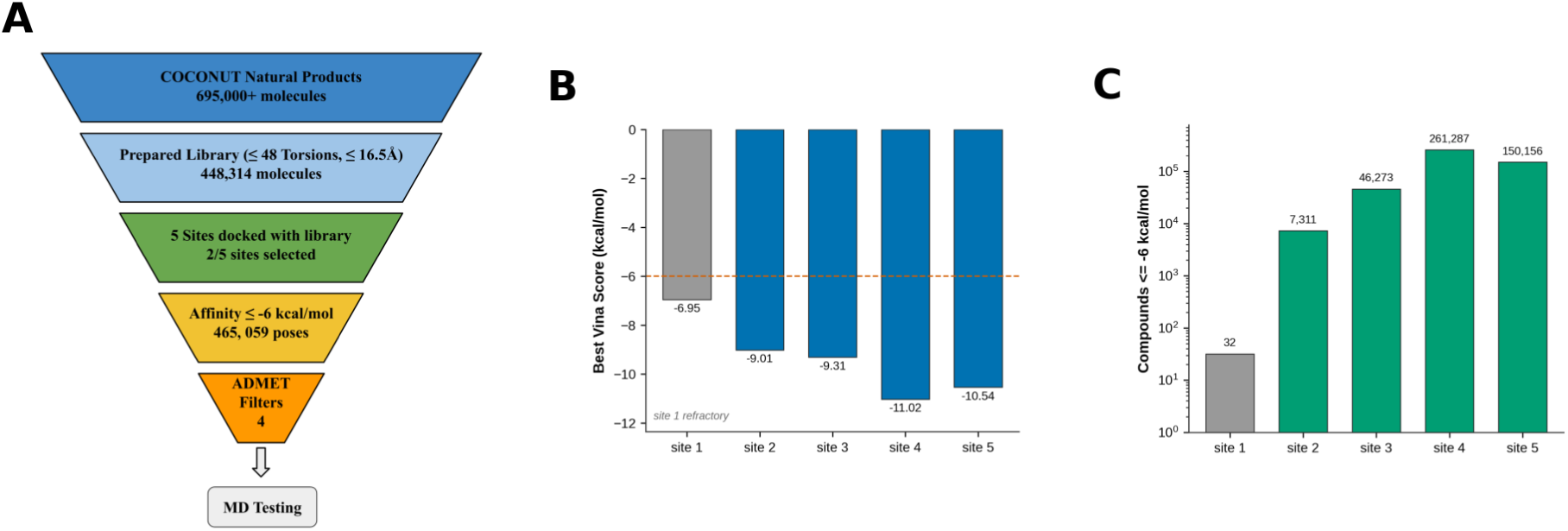
High-throughput docking of 448,314 COCONUT natural products across five hotspots. (A) Screening funnel from COCONUT source records. (B) Best Vina score per site. (C) Number of compounds scoring ≤ −6 kcal/mol per site (log scale).

**Table 2.** Docking Scores by Site. Uni-Dock/Vina score distributions for the 448,314 COCONUT compounds docked into each of the five hotspot-derived sites.

| Site | Box center (x, y, z; $\text{\AA}$) | Compounds scored | Best score (kcal/mol) | Compounds $\leq -6$ kcal/mol |
| --- | --- | --- | --- | --- |
| site01 | 78.7, 117.0, 40.8 | 425,475 | -6.95 | 32 |
| site02 | 89.6, 102.4, 31.8 | 407,300 | -9.01 | 7311 |
| site03 | 89.4, 108.2, 49.8 | 438,457 | -9.31 | 46,273 |
| site04 | 89.4, 118.3, 32.2 | 441,353 | -11.02 | 261,287 |
| site05 | 90.1, 112.5, 32.1 | 431,108 | -10.54 | 150,156 |

### 3.4 Diffusion-Based Design Yields Peptide Binders to the PC1–PC2 Interface, PC2 Regulatory Loop and Catalytic Region

To explore alternative modalities that better exploit the surfaces that resulted from cosolvent mapping, peptide binders were generated with BoltzGen based on three mechanistically distinct target regions (the PC1–PC2 subdomain interface, the PC2 regulatory loop and the catalytic region), with 20 designs per site. One design per region was chosen after filtration through short 30 ns simulations scored by per-residue RMSF, a subsequent 100 ns simulation, and MM-PBSA scoring of the three candidates per site with the lowest fluctuation.

The interface design (PC1–PC2; GRAPPGPPPPPRPPPTGRPGGVT, 23 residues) is proline-rich. The PC2-loop design (PC2L1; AVTRGSRWVPPRP, 13 residues) is shorter and combines basic and aromatic residues. The catalytic-region design (Triad; GERGPPGPRS, 10 residues) is the most compact of the three.

Co-folding each design with CAPN5 in Boltz-2, using the 6P3Q protease-core template with a pocket constraint over approximately 41 contact residues and 25 diffusion samples per peptide, returned high interface confidence for all three designs (Fig. 4B; Table 3). The catalytic-region and PC2-loop designs were both confidently and consistently placed (best-model ipTM 0.948 and 0.936). The interface design was less well determined (best-model ipTM 0.905).

**Figure 4.**
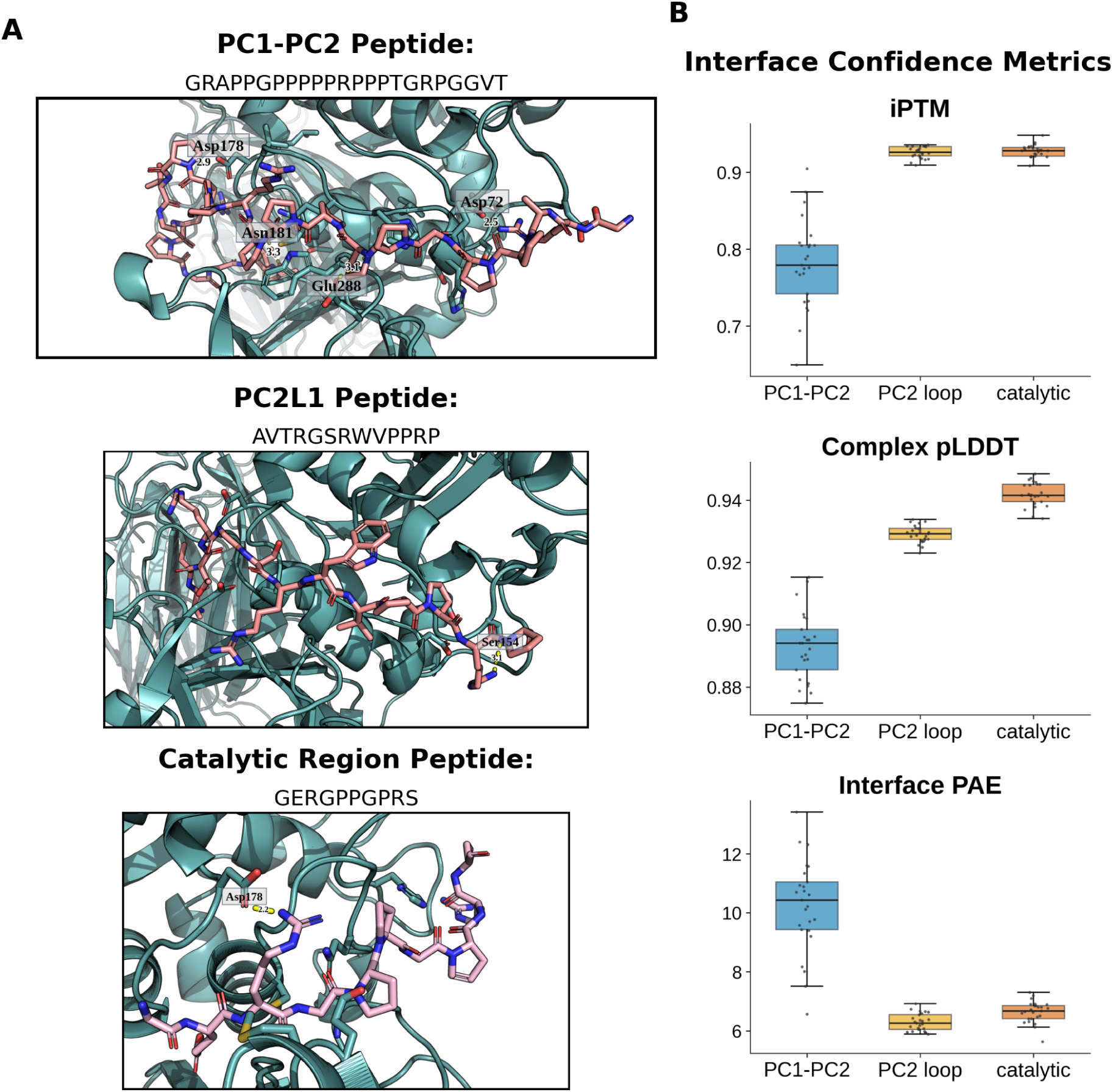
Diffusion-based peptide design and co-folding validation. (A) Boltz-2 co-folded complexes for the three retained designs. Only contacts at or under 3.5 Å are drawn, each labelled with the protein residue it reaches (boxed) and its donor–acceptor distance. All distances are in ångströms. (B) Interface confidence metrics (ipTM, complex pLDDT and interface PAE) across the 25 diffusion samples per design.

**Table 3.** Designed Peptide Binders. BoltzGen-designed peptide binders, Boltz-2 co-folding confidence over 25 diffusion samples per design, and interface geometry measured from the MD trajectories.

|  | PC1-PC2 | PC2L1 | Triad |
| --- | --- | --- | --- |
| Sequence | GRAPPGPPPP<br>RPPPTGRPGGVT | AVTRGSRWVPPRP | GERGPPGPRS |
| Length | 23 | 13 | 10 |
| ipTM (best) | 0.905 | 0.936 | 0.948 |
| ipTM (mean ± SD) | 0.782 ± 0.057 | 0.926 ± 0.007 | 0.927 ± 0.009 |
| Interface PAE ($\text{\AA}$) | 10.23 | 6.29 | 6.64 |
| Complex pLDDT | 0.893 | 0.929 | 0.942 |
| Buried SASA ($\text{\AA}^2$) | 2209 ± 279 | 1952 ± 44 | 1542 ± 45 |
| Peptide buried (%) | 45.3 | 56.5 | 62.8 |
| Most persistent salt bridge (occ.) | Arg12-Glu241<br>(43.8%) | Arg7-Glu241<br>(49.5%) | Glu2-Lys233<br>(67.2%) |

Interface geometry, however, favors the interface design which buries by far the largest interface (2209 ± 279 Å^2^ of surface, 1105 Å^2^ of interface area) but only 45.3% of the peptide itself, consistent with an extended proline-rich chain lying across a broad, shallow surface with substantial residual mobility. The catalytic-region design buries the least total surface (1542 ± 45 Å^2^) but the greatest fraction of itself (62.8%), consistent with a compact peptide seated in a defined pocket, and it forms the most persistent electrostatic contact of the set (Glu2–Lys233 salt bridge, 67.2% occupancy, mean lifetime 0.70 ns). The PC2-loop design is intermediate on both measures and forms the most persistent hydrogen bond (Arg155 N–H··· O Arg12, 76.3% occupancy).

The catalytic-region design anchors on Lys233, which is itself a contact of the catalytic hotspot, so that design engages the surface it was directed at. The interface and PC2-loop designs both anchor on the same residue, Glu241, at 43.8% and 49.5% occupancy; Glu241 appears in no hotspot contact set, and the region they occupy is the gating loop. These peptide binders produce a trend in which binding confidence was inversely proportional to the size of the binding interface.

### 3.5 MD and MM-PBSA Rank Binders Across Wild Type and ADNIV Variants

The top three peptides and four small molecules were simulated against wild-type CAPN5 and the four ADNIV variants for 100 ns and rescored by MM-PBSA with per-residue decomposition. Each of these 35 binder–background conditions was run in independent triplicate, so the 105 resulting trajectories carry a replicate spread for every cell of the matrix rather than a single number; error bars in Fig. 5 and the uncertainties in Table 4 are that between-replicate standard deviation. The runs showed consistent protease-core folding and per-residue C*α* RMSF concentrated on the gating loop and the grafted periphery, away from the catalytic core (Fig. 5A). All ligands were uniformly favorable, with peptides showing greater variation than small molecules.

**Figure 5.**
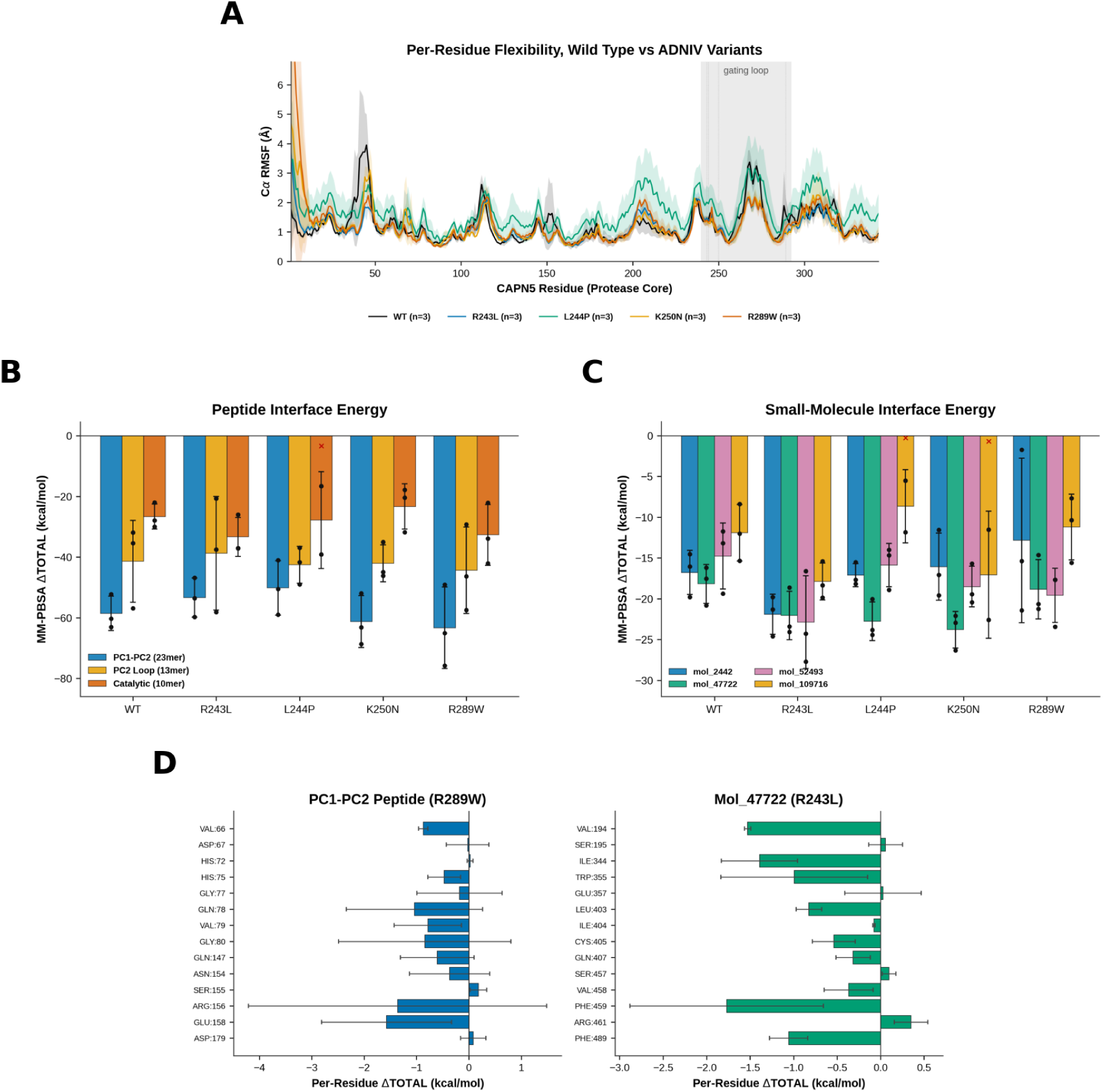
Molecular dynamics and MM-PBSA across wild type and ADNIV variants. Every panel is built from three independent 100 ns replicates per condition. (A) Per-residue C*α* RMSF over the protease core for wild-type CAPN5 and the four ADNIV variants, traces averaged over replicates with the replicate spread shaded, the gating loop shaded and the four variant positions marked. (B) MM-PBSA ΔTOTAL for the three peptide binders across all five backgrounds and (C) for the four small molecules. Red crosses mark the three replicates in which the ligand dissociated during the trajectory. (D) Per-residue energy decomposition for the strongest peptide (PC1–PC2 vs R289W) and the most consistent small molecule (Mol 47722 vs R243L).

**Table 4.**
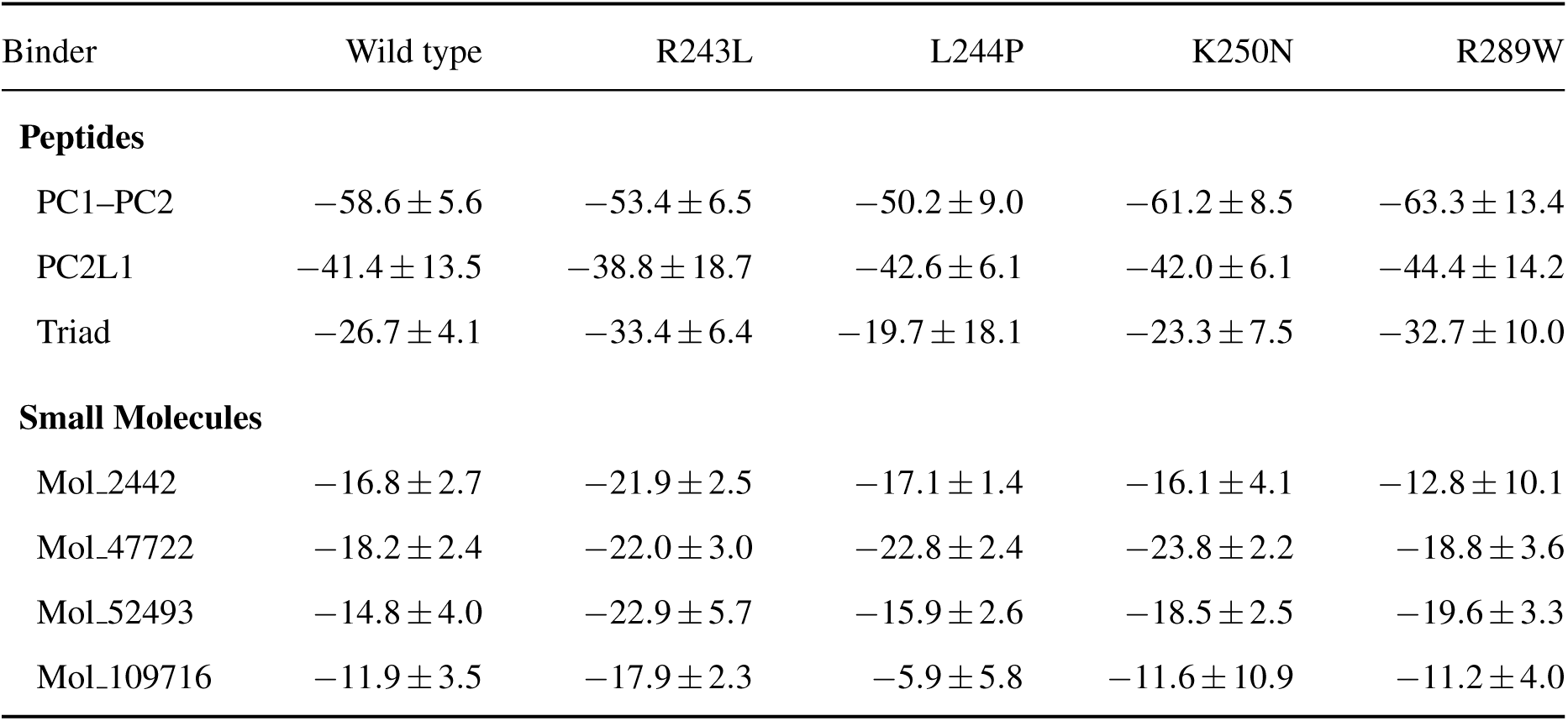
MM-PBSA Interface Energies. ΔTOTAL, kcal/mol, mean ± standard deviation *across independent replicates*, for each binder against wild-type CAPN5 and the four ADNIV variants.

In wild type, the interface design was strongest at ΔTOTAL −58.6 ± 5.6 kcal/mol, followed by the PC2-loop design at −41.4 ± 13.5 kcal/mol and the catalytic-region design at −26.7 ± kcal/mol (Fig. 5B). The same ordering held across all four variant backgrounds, with the interface design spanning −50.2 to −63.3 kcal/mol and the catalytic-region design −23.3 to −33.4 kcal/mol.

The small-molecule panel returned smaller interface energies, spanning −8.7 to −23.8 kcal/mol across the matrix (Fig. 5C). Their replicate standard deviations are 1.4–10.1 kcal/mol, overlapping the peptide panel’s relative spread rather than sitting well below it.

Each binder has a unique preferred background. The catalytic-region peptide is most favorable against R243L (−33.4 ± 6.4 kcal/mol against its own wild-type value of −26.7 ± 4.1 kcal/mol), whereas both longer peptides peak against R289W; among the small molecules, Mol 52493 and Mol 109716 peak against R243L and Mol 47722 against K250N. The differences are 3 to 8 kcal/mol against a median replicate standard deviation of 5.0 kcal/mol, so no single assignment is established by three replicates; taken together they suggest matching a binder to a patient’s substitution, and identify which pairings a larger replicate set should test first.

## 4 Conclusions

We assembled and executed an *in silico* pipeline to produce candidate CAPN5 binders for exploring inhibition for ADNIV. We grafted an AlphaFold model onto the 6P3Q protease-core crystal structure and mapped the resulting full-length construct using phenol cosolvent MD. This analysis defined the druggable pockets of CAPN5 and identified five potential binding regions. We docked 448,314 COCONUT natural products against all five regions and used diffusion-based methods to design peptides. MM-PBSA analysis on peptides show increasing affinity with peptide length and buried interface area. A 105-simulation matrix spanning wild type and the R243L, L244P, K250N and R289W variants show binder background preferences. Confirming any individual pairing would need a larger replicate set, as the differences are smaller than three replicates show. These results characterize CAPN5 as a logical target and identify the catalytic cleft and the adjoining groove best for inhibition.

## CRediT Authorship Contribution Statement

**Surya Charan Garimella:** Methodology, Software, Investigation, Formal analysis, Data curation, Visualization, Writing (original draft). **Yash Bhargava:** Conceptualization, Methodology, Software, Investigation, Formal analysis, Supervision, Writing (review & editing). Both authors read and approved the final manuscript.

## Declaration of Competing Interest

The authors declare that they have no known competing interests that influenced the work reported in this paper.

## Data Availability

The original contributions presented in the study are included in the article and its Supplementary Material. Further inquiries can be directed to the corresponding author. Molecular-dynamics trajectories and the complete pipeline archive are available from the corresponding author on request.

## Notes

### Competing Interest Statement

The authors have declared no competing interest.

