## Supplementary Figures and Tables for "*In Silico* Targeting of Calpain-5 in Autosomal Dominant Neovascular Inflammatory Vitreoretinopathy (ADNIV) with Peptides and Natural Products"

Surya Charan Garimella Yash Bhargava

Bhargava Systems Research Inc., Carmel, IN, USA

### Supplementary Figures

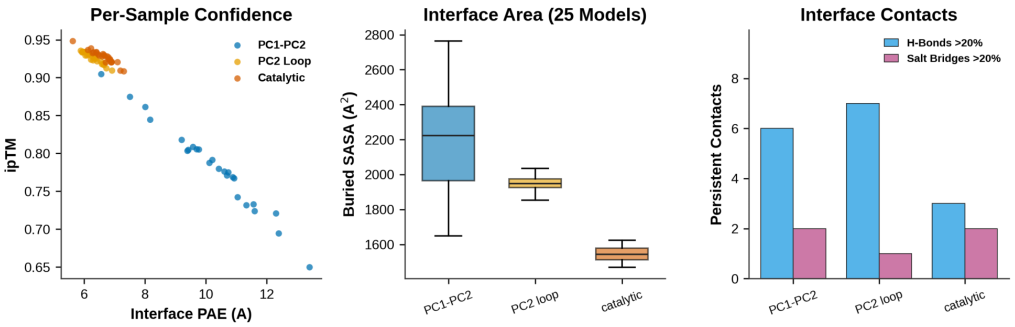


**Figure S1. Boltz-2 co-folding interface detail.** (A) Per-sample ipTM versus mean interface predicted aligned error for the 25 diffusion samples of each design. (B) Buried solvent-accessible surface area across the 25 models per design. (C) Counts of hydrogen bonds and salt bridges occupied in more than 20% of the co-folded models.

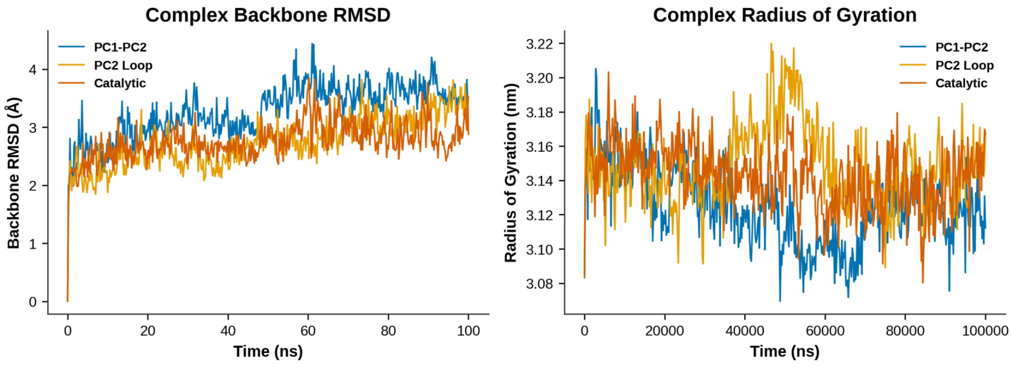


**Figure S2. Molecular-dynamics stability of the peptide complexes.** Complex backbone RMSD (left) and radius of gyration (right) over the 100 ns wild-type trajectories for the three peptide designs.

### Supplementary Tables

**Table S1. Full Hotspot Cluster Metrics.** DBSCAN cluster metrics for the five phenol hotspots on CAPN5 (500 ns cosolvent simulation), extending Table 1 of the main text with cluster radius and nearest-protein distance.

| Rank | Phenol obs. | Frame occ. (%) | Max-cell enrich. (×) | Mean-cell enrich. (×) | Cluster radius (Å) | Nearest protein heavy atom (Å) |
|---|---|---|---|---|---|---|
| 1 | 3307 | 66.1 | 4066 | 1173 | 0.77 | 1.88 |
| 2 | 2359 | 47.2 | 1761 | 837 | 0.86 | 0.76 |
| 3 | 2208 | 44.2 | 1634 | 783 | 0.79 | 1.81 |
| 4 | 1680 | 33.6 | 1346 | 813 | 0.75 | 1.67 |
| 5 | 1465 | 29.3 | 1543 | 780 | 0.84 | 2.03 |

**Table S2. Per-Hotspot Residue Contacts.** Residue-contact fractions (top eight residues per hotspot), the fraction of that cluster's probe observations that contact each residue.

| Hotspot rank | Residue | Contact count | Contact fraction |
|---|---|---|---|
| 1 | ARG283 | 3307 | 1.000 |
| 1 | SER69 | 3307 | 1.000 |
| 1 | SER88 | 3307 | 1.000 |
| 1 | HIS71 | 3307 | 1.000 |
| 1 | PRO285 | 3304 | 0.999 |
| 1 | ASP72 | 3303 | 0.999 |
| 1 | LEU73 | 3299 | 0.998 |
| 1 | SER92 | 3279 | 0.992 |
| 2 | LYS339 | 2359 | 1.000 |
| 2 | ILE197 | 2358 | 1.000 |
| 2 | ILE337 | 2356 | 0.999 |
| 2 | LEU214 | 2349 | 0.996 |
| 2 | MET218 | 2341 | 0.992 |
| 2 | ARG217 | 2339 | 0.992 |
| 2 | GLU195 | 2336 | 0.990 |
| 2 | ILE338 | 2294 | 0.972 |
| 3 | LYS99 | 2207 | 1.000 |
| 3 | THR190 | 2205 | 0.999 |
| 3 | LEU96 | 2196 | 0.995 |
| 3 | PHE189 | 2192 | 0.993 |

*Table S2 continued*

| Hotspot rank | Residue | Contact count | Contact fraction |
|---|---|---|---|
| 3 | TRP131 | 2186 | 0.990 |
| 3 | VAL100 | 1934 | 0.876 |
| 3 | LEU90 | 1747 | 0.791 |
| 3 | HIS124 | 1578 | 0.715 |
| 4 | GLY251 | 1679 | 0.999 |
| 4 | ALA253 | 1679 | 0.999 |
| 4 | CYS81 | 1677 | 0.998 |
| 4 | HIS252 | 1667 | 0.992 |
| 4 | SER231 | 1628 | 0.969 |
| 4 | ALA85 | 1549 | 0.922 |
| 4 | TRP82 | 1544 | 0.919 |
| 4 | TRP286 | 1521 | 0.905 |
| 5 | ILE338 | 1465 | 1.000 |
| 5 | SER231 | 1465 | 1.000 |
| 5 | ASP336 | 1464 | 0.999 |
| 5 | THR182 | 1463 | 0.999 |
| 5 | SER229 | 1463 | 0.999 |
| 5 | ILE337 | 1461 | 0.997 |
| 5 | ALA230 | 1454 | 0.992 |
| 5 | ALA253 | 1312 | 0.896 |

**Table S3. Per-Replicate MM-PBSA Results.** Complete MM-PBSA results across the 105-replicate matrix (7 binders $\times$ 5 backgrounds $\times$ 3 replicates) for 100 ns at 310 K.

| Target | Variant | Rep. | PB ΔTOTAL (kcal/mol) | Within-traj. SD |
|---|---|---|---|---|
| PC1–PC2 | WT | rep1 | −52.28 | 14.79 |
| PC1–PC2 | WT | rep2 | −63.08 | 8.69 |
| PC1–PC2 | WT | rep3 | −60.32 | 8.93 |
| PC1–PC2 | R243L | rep1 | −53.58 | 10.82 |
| PC1–PC2 | R243L | rep2 | −59.71 | 9.67 |
| PC1–PC2 | R243L | rep3 | −46.79 | 9.01 |
| PC1–PC2 | L244P | rep1 | −50.51 | 15.64 |
| PC1–PC2 | L244P | rep2 | −58.99 | 6.92 |

*Table S3 continued*

| Target | Variant | Rep. | PB ΔTOTAL (kcal/mol) | Within-traj. SD |
|---|---|---|---|---|
| PC1–PC2 | L244P | rep3 | −41.01 | 13.95 |
| PC1–PC2 | K250N | rep1 | −68.68 | 14.84 |
| PC1–PC2 | K250N | rep2 | −63.15 | 8.10 |
| PC1–PC2 | K250N | rep3 | −51.92 | 9.01 |
| PC1–PC2 | R289W | rep1 | −75.79 | 7.97 |
| PC1–PC2 | R289W | rep2 | −65.02 | 9.40 |
| PC1–PC2 | R289W | rep3 | −49.11 | 8.58 |
| PC2L1 | WT | rep1 | −35.44 | 18.00 |
| PC2L1 | WT | rep2 | −31.90 | 7.61 |
| PC2L1 | WT | rep3 | −56.85 | 8.32 |
| PC2L1 | R243L | rep1 | −37.49 | 22.76 |
| PC2L1 | R243L | rep2 | −20.74 | 10.63 |
| PC2L1 | R243L | rep3 | −58.12 | 10.13 |
| PC2L1 | L244P | rep1 | −36.98 | 4.28 |
| PC2L1 | L244P | rep2 | −49.05 | 7.40 |
| PC2L1 | L244P | rep3 | −41.69 | 8.97 |
| PC2L1 | K250N | rep1 | −44.94 | 15.59 |
| PC2L1 | K250N | rep2 | −35.08 | 6.93 |
| PC2L1 | K250N | rep3 | −46.13 | 8.64 |
| PC2L1 | R289W | rep1 | −46.33 | 12.11 |
| PC2L1 | R289W | rep2 | −57.46 | 11.93 |
| PC2L1 | R289W | rep3 | −29.26 | 10.47 |
| Triad | WT | rep1 | −27.94 | 3.85 |
| Triad | WT | rep2 | −22.05 | 4.59 |
| Triad | WT | rep3 | −30.04 | 10.49 |
| Triad | R243L | rep1 | −37.11 | 13.98 |
| Triad | R243L | rep2 | −37.04 | 6.73 |
| Triad | R243L | rep3 | −25.98 | 7.85 |
| Triad | L244P | rep1 | −16.58 | 24.64 |
| Triad | L244P | rep2 | −39.12 | 12.21 |
| Triad | L244P | rep3 | −3.37 [†] | 4.03 |
| Triad | K250N | rep1 | −20.39 | 6.49 |
| Triad | K250N | rep2 | −17.81 | 8.11 |
| Triad | K250N | rep3 | −31.82 | 12.14 |

*Table S3 continued*

| Target | Variant | Rep. | PB ΔTOTAL (kcal/mol) | Within-traj. SD |
|---|---|---|---|---|
| Triad | R289W | rep1 | −22.17 | 9.16 |
| Triad | R289W | rep2 | −42.05 | 6.72 |
| Triad | R289W | rep3 | −33.91 | 5.73 |
| Mol_2442 | WT | rep1 | −19.78 | 3.04 |
| Mol_2442 | WT | rep2 | −14.54 | 4.87 |
| Mol_2442 | WT | rep3 | −16.03 | 4.26 |
| Mol_2442 | R243L | rep1 | −19.78 | 1.03 |
| Mol_2442 | R243L | rep2 | −24.59 | 3.85 |
| Mol_2442 | R243L | rep3 | −21.33 | 3.48 |
| Mol_2442 | L244P | rep1 | −15.51 | 0.72 |
| Mol_2442 | L244P | rep2 | −18.18 | 3.71 |
| Mol_2442 | L244P | rep3 | −17.66 | 3.35 |
| Mol_2442 | K250N | rep1 | −19.57 | 0.94 |
| Mol_2442 | K250N | rep2 | −11.55 | 4.67 |
| Mol_2442 | K250N | rep3 | −17.08 | 3.42 |
| Mol_2442 | R289W | rep1 | −1.72 | 1.58 |
| Mol_2442 | R289W | rep2 | −21.43 | 4.02 |
| Mol_2442 | R289W | rep3 | −15.38 | 2.77 |
| Mol_47722 | WT | rep1 | −17.54 | 5.41 |
| Mol_47722 | WT | rep2 | −20.81 | 3.51 |
| Mol_47722 | WT | rep3 | −16.19 | 2.96 |
| Mol_47722 | R243L | rep1 | −24.08 | 0.82 |
| Mol_47722 | R243L | rep2 | −23.41 | 2.11 |
| Mol_47722 | R243L | rep3 | −18.64 | 4.04 |
| Mol_47722 | L244P | rep1 | −23.84 | 1.05 |
| Mol_47722 | L244P | rep2 | −20.03 | 4.22 |
| Mol_47722 | L244P | rep3 | −24.42 | 2.52 |
| Mol_47722 | K250N | rep1 | −22.95 | 1.16 |
| Mol_47722 | K250N | rep2 | −26.34 | 3.33 |
| Mol_47722 | K250N | rep3 | −22.09 | 3.78 |
| Mol_47722 | R289W | rep1 | −14.64 | 1.38 |
| Mol_47722 | R289W | rep2 | −20.64 | 2.76 |
| Mol_47722 | R289W | rep3 | −21.24 | 3.43 |
| Mol_52493 | WT | rep1 | −11.75 | 5.45 |

*Table S3 continued*

| Target | Variant | Rep. | PB ΔTOTAL (kcal/mol) | Within-traj. SD |
|---|---|---|---|---|
| Mol_52493 | WT | rep2 | −19.37 | 3.42 |
| Mol_52493 | WT | rep3 | −13.17 | 3.15 |
| Mol_52493 | R243L | rep1 | −24.28 | 2.79 |
| Mol_52493 | R243L | rep2 | −27.72 | 2.96 |
| Mol_52493 | R243L | rep3 | −16.62 | 5.21 |
| Mol_52493 | L244P | rep1 | −14.68 | 1.30 |
| Mol_52493 | L244P | rep2 | −18.90 | 3.82 |
| Mol_52493 | L244P | rep3 | −14.00 | 3.84 |
| Mol_52493 | K250N | rep1 | −19.43 | 2.44 |
| Mol_52493 | K250N | rep2 | −20.42 | 4.38 |
| Mol_52493 | K250N | rep3 | −15.68 | 2.72 |
| Mol_52493 | R289W | rep1 | −17.69 | 2.11 |
| Mol_52493 | R289W | rep2 | −17.66 | 3.13 |
| Mol_52493 | R289W | rep3 | −23.42 | 4.74 |
| Mol_109716 | WT | rep1 | −15.33 | 3.27 |
| Mol_109716 | WT | rep2 | −8.39 | 3.61 |
| Mol_109716 | WT | rep3 | −12.01 | 4.27 |
| Mol_109716 | R243L | rep1 | −19.88 | 2.06 |
| Mol_109716 | R243L | rep2 | −15.42 | 3.64 |
| Mol_109716 | R243L | rep3 | −18.35 | 1.91 |
| Mol_109716 | L244P | rep1 | −5.50 | 2.55 |
| Mol_109716 | L244P | rep2 | −0.25 † | 1.34 |
| Mol_109716 | L244P | rep3 | −11.84 | 4.41 |
| Mol_109716 | K250N | rep1 | −22.57 | 1.17 |
| Mol_109716 | K250N | rep2 | −0.67 † | 1.92 |
| Mol_109716 | K250N | rep3 | −11.54 | 2.60 |
| Mol_109716 | R289W | rep1 | −7.68 | 2.52 |
| Mol_109716 | R289W | rep2 | −10.34 | 6.01 |
| Mol_109716 | R289W | rep3 | −15.60 | 3.07 |
